# Reconciling Two Sides: Novel Analytical Statistics for Lesion Network Mapping

**DOI:** 10.64898/2026.09.09.750470

**Authors:** Azamat Yeldesbay, Silvia Daun

## Abstract

Recent studies have reached conflicting conclusions about the statistical significance of lesion network mapping (LNM), especially regarding the impact of regional node strength on lesion-associated network detection. Here, we present a statistical framework that clarifies the origin of these apparently conflicting findings. Based on the simplified formulation of LNM introduced by van den Heuvel et al. (2026), we propose alternative, permutation-based null models corresponding to the sensitivity and specificity tests used in standard LNM analysis. We derive analytical expressions for both the proposed null models and the node-strength-constrained approach recently introduced by Zalesky and Cash (2026), thereby providing a computationally efficient alternative to permutation-based significance testing. Using synthetic lesion and connectivity matrices, as well as clinical datasets, we compare the outcome of the proposed null models with the standard LNM analysis and demonstrate how the choice of null models influences statistical inference and the resulting lesion network maps. Our framework shows that the apparent disagreements in the recent LNM literature can be understood as a consequence of different statistical null models rather than to conflicting conclusions about LNM itself.

## 1 Introduction

Lesion Network Mapping (LNM) is a widely used framework for investigating the relationship between focal brain lesions and neurological or psychiatric symptoms. It leverages normative functional connectivity data. Over the past decade, this method has been applied to a wide variety of clinical conditions, demonstrating its potential to identify distributed brain networks associated with lesion-induced deficits [1, 2].

Recently, [3] proposed a simplified formulation of LNM, arguing that much of the observed results could be explained by regional node strength. These findings have sparked a substantial discussion about the statistical interpretation of LNM results and the question of whether the reported associations arise from lesion-specific effects or from the underlying network topology. Several studies [4–6] have challenged the conclusions of [3], arguing that significant lesion-network relationships remain after appropriate statistical controls and highlighting the limitations of the simplified analysis. Other authors have proposed alternative statistical frameworks for assessing LNM significance [6–8]. For instance, one such framework proposed a null model based on permutations constrained to regions with similar node strength, thereby attempting to control for this important network property [7]. Moreover, permutation-based improvements as suggested in several studies [6, 7] had already been adopted by the LNM community [9, 10]. However, a direct comparison between the performances of permutation-based and standard LNM analyses is still lacking in the literature.

In this work, we present a novel, analytically formulated statistical framework that reconciles the seemingly conflicting conclusions of recent LNM studies. To this end, we introduce alternative, permutation-based null distributions for statistical testing at the level of individual brain regions that correspond to the sensitivity and specificity tests used in the standard LNM analysis. The proposed null models are derived from the simplified LNM formulation introduced by [3], and are designed to reduce the strong dependence of LNM on regional node strength while improving lesion-associated network detection. We derive analytical expressions for the proposed null distributions and the node-strength-constrained null distribution introduced by [7]. We evaluate both approaches using the synthetic connectivity simulations proposed by [7] and with clinical lesion datasets, including those distributed with the implementation of [3]. Furthermore, we compare the complete LNM analysis pipeline, including sensitivity, specificity, and conjunction map analyses, between the proposed method and the standard LNM framework. Finally, we demonstrate how the proposed statistical framework helps explain the apparently conflicting conclusions regarding the statistical significance of LNM reported in the recent literature.

This paper is organized as follows. First, we introduce the standard LNM as originally introduced in [11] (i.e., without permutation). Then, we define the mathematical formulation of LNM and derive the analytical forms of the proposed null models for the tests in the standard LNM analysis. Next, we present the results of the synthetic validation simulations and the analyses performed on clinical datasets. Finally, we compare the performance of the different statistical approaches and discuss their implications for interpreting and applying lesion network mapping in the future.

## 2 Standard Lesion Network Mapping Analysis

The standard lesion network mapping analysis consists of the following tests [1, 4].

### Sensitivity Test

In the original analysis, a sensitivity map was calculated by thresholding and binarizing seed maps from each region and summing them up to create a frequency map. Some groups, however, have replaced this step by calculating the t-values of the Fisher r-to-z transformed correlation values for each region. Thus, the null hypothesis has a distribution centered around zero. In the simplified formulation of LNM ([3], Eq. (1)), this is equivalent to finding the significance of the mean of the columns of the matrix **LNM** against zero.

The use of a t-test was deliberately avoided in the LNM analysis, since it assumes zero-centricity [11, 12]. Here, however, we compare our results with those of the one-sample t-test, because seed-map thresholding is not possible in a matrix formulation: **C** already contains averaged r-values across subjects of the normative connectome dataset.

### Specificity Test

The specificity test aims to identify the regions specific to a given clinical symptom. This test is almost always included in the LNM analysis [4], since the specificity map itself is not controlled against any baseline. In the standard LNM analysis, there are two groups of measurements: one group includes lesions that cause a given clinical symptom, and the other group includes lesions that cause no or different clinical symptoms. The test was originally performed as a region-wise two-sample t-test. However, recent studies have applied nonparametric analysis instead, which carries out a permutation of the clinical scores of the subjects in the specificity test [9, 10]. Here, we compare our results with those of the two-sample t-test to demonstrate the differences with the permutation analysis, thereby addressing the original critique in [3].

### Conjunction Map

The conjunction map combines the results of the sensitivity and specificity tests. The regions that are significant in both tests are selected.

## 3 Mathematical Formulation of the Proposed Statistical Framework

### Mathematical Formulation of LNM

In their recent work [3], van den Heuvel and his colleagues presented a simplified formulation of LNM as a matrix multiplication of the lesion matrix and the connectivity matrix:

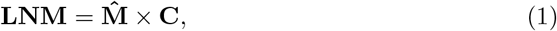

where **C** is an *R × R* group-level functional connectivity matrix averaged across subjects in a normative resting-state functional connectivity dataset, and 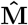 is an *S × R* lesion matrix, with elements 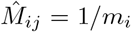 if the lesion covers the region *j* for subject *i* or 0 otherwise; *m*_*i*_ is the size of the lesion for subject *i, S* and *R* are the number of subjects and brain regions, respectively. The resulting matrix, **LNM**, of size *S × R* represents the correlation between brain regions and lesion sides for all subjects. The lesion matrix 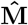 is derived from an *S × R* binary lesion matrix, **M**, via row-normalization (see Supplementary Section S1 for details on the definitions).

Based on the simplified formulation Eq. (1) and its modification [7], we define LNM to quantify the average functional connectivity between the lesion and each brain region across subjects, as well as to identify regions whose connectivity to the lesion network is associated with the clinical symptom:

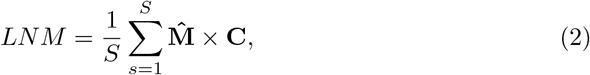

and redefine this expression as

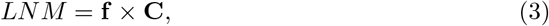

where 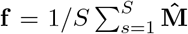 is a vector of the mean frequencies of regions to be affected by lesions across all subjects. This is equivalent to the lesion overlap map (see Supplementary Section S1 for details on the definitions).

### Null Model for Sensitivity Test

We derive a null model for the sensitivity test based on permutation by asking if the observed value of *LNM*_*k*_ for a region *k* is significantly larger than what would be produced by random lesion-region assignment (see Supplementary Section S2 for details on the derivation). In the expressions Eqs. (2) and (3), the connectivity matrix is fixed for a given connectivity dataset. Thus, the random variables are the lesion matrix 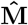 and the frequency vector **f** . This means that the null distribution for the sensitivity test is constructed by permuting the columns of the lesion matrix 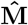 (see Fig. 1 **a**), which is equivalent to shuffling the lesion overlap map across the brain regions (see Fig. 1 **b**). Deriving the null distribution for LNM yielded an analytic expression for the expected value of 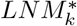^1^ for region *k*:

**Fig. 1.**
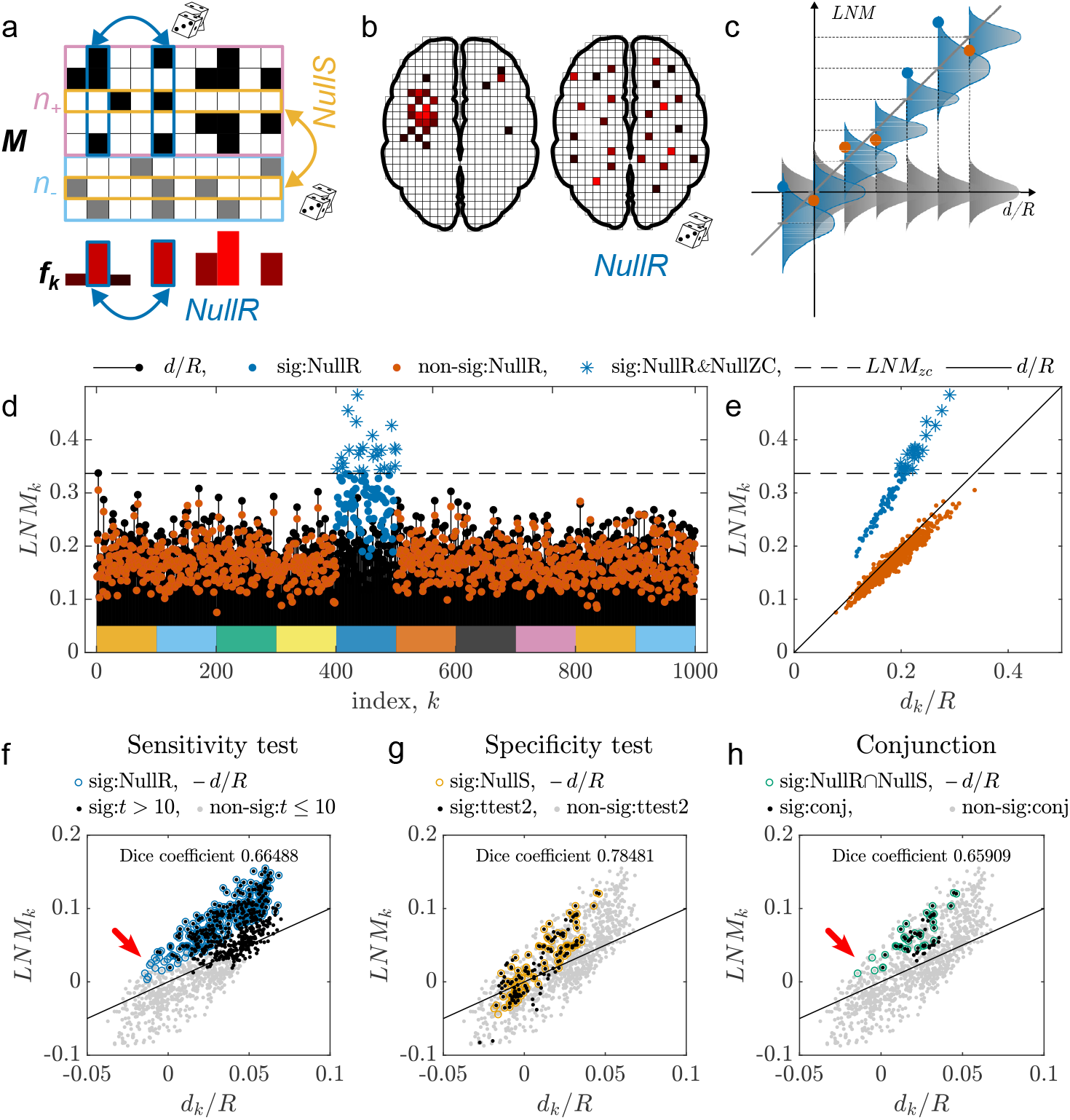
Explanation of the alternative statistical method for LNM and its validation using synthetic and clinical data. **a**, The null models are based on the permutation of the lesion matrix **M** in two directions: *NullR* uniform random permutation of the regions (blue), *NullS* uniform random permutation of the rows (light orange), which are divided into two groups: upper *n*_+_ rows with (black squares surrounded by pink rectangle) and lower *n*_−_ rows without (gray squares surrounded by light blue rectangle) symptoms. **b**, *NullR* permutation is equivalent to permuting the mean frequency of occurrence of a lesion in a given region or permuting the lesion overlap map. **c**, the expected LNM value for a region depends on its mean node strength (gray diagonal solid line). *NullR* distribution is approximated as normal distribution for each region separately (blue distributions). In the standard LNM sensitivity test, a zero-centered null distribution is used (gray distributions). **d, e**, show the results of the validation with synthetic connectivity and lesion matrices: **d** The regions of the target module (blue dots and stars) appear statistically significant in *NullR* (*p <* 0.05, Bonferroni corrected), whereas the other module regions do not (orange dots). Blue stars denote regions significant according to the Zalesky & Cash null model (*NullZC*); the threshold (*LNM*_*zc*_) is denoted by the dashed line. The value of the lesion distribution parameter is *α* = 0.8. The black dots and solid lines show the expected values. **e**, *LNM* values plotted against region’s averaged node degree *d/R*. Strongly interconnected regions with higher lesion occurrence have *LNM* values (blue dots and stars) that significantly deviate from the expected value (black solid line). **f**,**g**,**h** show the results of the validation using a clinical dataset. Each panel depicts the mean *LNM*_*k*_ values for individual regions *k* on the (*LNM, d/R*) plane, with each dot representing one region. **f**, Results of the sensitivity test. Black and gray dots denote the regions significant (with *t >* 10 against zero mean value) and not significant in the standard sensitivity test. The blue circles denote the regions significant in *NullR* and the red arrow points to the ones with lower node strength. The results of both tests highly overlap (dice coefficient of 0.66488). **g**, Results of specificity test. Yellow circles mark the regions statistically significant according to *NullS*, and black and gray dots denote the significant and non-significant ones according to region-specific two sample t-test used in the standard specificity test. The results of the two tests highly overlap (dice coefficient of 0.78481) **h**, Results of the conjunction map. Green circles denote the regions significant in *NullR* and *NullS*, red arrow points to the ones with lower node strength, and black and gray dots denote the regions identified in the standard conjunction test. The overlap is also high (dice coefficient of 0.65909).

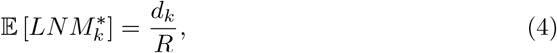

where 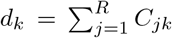 is the node strength of region *k*. Additionally, we found an analytic expression for the variance:

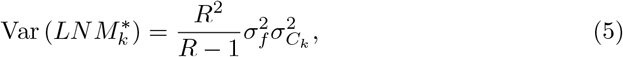

where 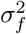 is the variance of **f** across permutation, and 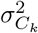 is the variance of column *k* of the matrix **C** (see Supplementary Section S2 for details on the derivation and S4 for the verification of the analytic expressions using permutation).

These results imply that the significance of *LNM* should not be assessed relative to zero, but relative to an expected value that depends linearly on the region’s average node strength. This supports the conclusion of van den Heuvel and colleagues [3] that LNM is biased by node strength. Thus, statistical testing must be performed region-wise against the corresponding null distribution rather than against a zero-mean null hypothesis (see Fig. 1 **c**).

In Eq. (5), the variance of the connectivity matrix **C** is fixed for a given dataset. Therefore, the variance of the LNM values is determined *exclusively* by the variance of the lesion frequencies **f**, which is defined by the lesion matrix 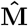 and reflects the distribution of lesions across subjects and regions. This allows for faster estimation of the variance than permutation does.

We refer to this model as *NullR* or column permutation because the permutation is performed across regions.

### Null Model for Specificity Test

The specificity test compares the mean *LNM* of two groups for every region *k*: one group has the given clinical symptom (+), while the other does not (−). The null model for the specificity test aims to determine the regions for which the difference in means between the two groups, 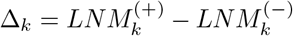, is significantly larger than what would be produced by randomly splitting the total number of subjects *S* = *n*_+_ + *n*_−_ into two groups of sizes *n*_+_ and *n*_−_. We find the null distribution by permuting the rows of a pooled lesion matrix 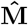 and selecting the first *n*_+_ rows as the group with the clinical symptom and the last *n*_−_ rows as the group without it (see Fig. 1 **a**).

This row-split permutation implies that the expected value of 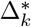 is zero for all regions:

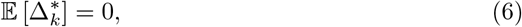

and the analytical expression for the variance is

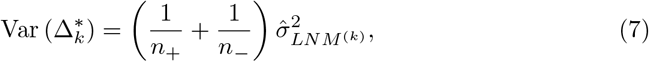

where 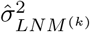 is the sample variance (over *S* − 1) of the *k*-th column of the **LNM** matrix calculated using the pooled lesion matrix (see Supplementary Section S2 for details on the derivation and S4 for the validation of the analytic expressions using permutation). The formulation of this null model is similar to the permutation analysis used in [9, 10], where permutation of the clinical scores of the subjects was performed.

In the following text, we refer to this null model as *NullS* or row-split permutation model.

### Conjunction Map

As in the standard LNM analysis, the conjunction map identifies regions that are significant in both the *NullR* and *NullS* tests.

### Zalesky & Cash Null Model

We also derived analytic expressions for the null model suggested by [7]. This model is based on permuting the lesions per subject within the “neighbors” of a region only. These neighbors are defined as the regions with the 20% closest node strengths in absolute value. The null distribution is built by finding the largest *LNM* value in each permutation. A region is considered significant in the sensitivity test if it exceeds the 95th percentile threshold of the null distribution (see Supplementary Section S3 for the analytical expressions and derivation details and Section S4 for the verification of the analytical formulae by the corresponding permutation). In the text, we refer to this null model as *NullZC*, and denote the 95th percentile threshold as *LNM*_*zc*_.

## 4 Validation of Null Models with Synthetic Connectivity and Lesion Matrices

To demonstrate the importance of performing sensitivity analyses as implemented in the *NullR* model, we used the synthetic connectivity and lesion matrix simulation framework presented by Zalesky and Cash [7] (see Supplementary Section S5 for details). In this test, 1000 regions were divided into 10 equal-sized modules, and a log-normal connectivity matrix was randomly generated with stronger region-to-region connections within modules than between modules. The parameter *α* regulates the probability of a lesion occurring in a target module: For *α* = 0, lesions are randomly distributed across the brain, whereas for *α* = 1, all lesions occur within the regions of the target module. Panel **d** of Fig. 1 shows the results for *α* = 0.8. The *NullR* test identifies all regions of the target module (blue circles and stars) as significant after Bonferroni correction (*p <* 0.05). In contrast, the *NullZC* model identifies the target module regions as statistically significant above the *LNM*_*zc*_ threshold (blue stars above the dashed line). Plotting the *LNM* values against *d/R* reveals a clear separation between the target module regions and the other regions (see Fig. 1 **e**). The *NullR* model can detect the target module regions even for smaller values of *α*. Furthermore, increasing *α* and/or the difference between intra-module and inter-module connectivity results in greater deviation of LNM values for the target module regions from the expected value (see Supplementary Section S5 for results with different parameter values).

## 5 Validation on a Clinical Dataset

We used a calculated, averaged connectivity matrix from the GSP1000 dataset [13] mapped to the Yeo-Schaefer1000 cortical [14] and Melbourne54 [15] sub-cortical regions. This matrix is available in the GitHub repository [16] of van den Heuvel and colleagues [3]. We used the Aphasia Recovery Cohort (ARC) [17] as a clinical dataset, the lesion matrix of which is also available in the GitHub repositories [16] and [18].

We validated the *NullR* and *NullS* models using the standard LNM analysis pipeline, which includes sensitivity and specificity tests and a conjunction map [4].

### Sensitivity Test

We compared the results of the standard sensitivity test with those of *NullR* (see Fig. 1 **f**). Unlike the synthetic case, real datasets do not have clear, separate regions on the (*LNM, d/R*) plane. Instead, they form a cloud with the first principal component (i.e., the direction with the largest variation in the data) tilted towards, but not aligned with, the diagonal line (i.e., the expected value). The angle between the first principal component and the diagonal line indicates an additional correlation between the occurrence of a lesion in a region and its node strength. This correlation is caused by the properties of the lesion matrix rather than the connectivity matrix (see Supplementary Section S6.4). However, due to the tilted shape, there is a large overlap between the two tests (dice coefficient is around 0.67 ^2^). We tested *LNM* values in other clinical cases and found that the shape of the LNM cloud varies and that the tilted shape is present in several cases (see Supplementary Section Fig. S14).

### Specificity Test

We identified groups with and without aphasia symptoms from the aphasia recovery cohort using a threshold of 93.5 for the clinical score of the Western Aphasia Battery - Aphasia Quotient (WAB-AQ). This resulted in the dataset being split into groups of *n*_+_ = 188 and *n*_−_ = 30 subjects.

The outcome of comparing *NullS* with the standard specificity test, i.e., the region-wise two-sample t-test (*ttest2*), is shown in Fig. 1 **g**. As can be seen, the regions that are significant in both specificity tests do not exhibit a bias towards higher node strength, which is a critical finding in relation to the original critique [3].

There is also a high overlap between the test results (dice coefficient of 0.78). However, a detailed comparison of the test results shows that *ttest2* can be strongly influenced by individual outlying observations for small sample sizes (see Supplementary Fig. S13). In contrast, *NullS* permutation testing is less sensitive to such observations. Furthermore, the two groups have the same expected mean *LNM* value, i.e., the averaged node strength of a region, which is independent of the sample sizes *n*_+_ and *n*_−_.

### Conjunction Map

Fig. 1 **h** shows the regions selected by the conjunction map in the standard LNM analysis as well as those that passed the *NullR* and *NullS* tests. Due to the specificity test results, the significant regions in both conjunction maps have intermediate and low node strength values and are not biased towards high values. The final overlap between the results of the standard LNM analysis and the statistical framework presented here is high (dice coefficient of 0.66). However, the *NullR* and *NullS* tests detect a set of regions with a wider range of node strength values than the standard conjunction map. The *NullR* test detects additional regions with lower node strength (red arrow in Fig. 1 **f**), some of which survive the specificity test and are thus present in the conjunction map (red arrow in Fig. 1 **h**).

## 6 Conclusions

The question of statistical significance of lesion network mapping is addressed by considering appropriate null models [7, 19]. We developed null models involving permutations of the columns and rows of the lesion matrix corresponding to LNM sensitivity and specificity tests, and derived analytical expressions for their corresponding null distributions.

The analytical expression for the *NullR* distribution corresponding to the sensitivity test reveals that its expected value is equal to the average node strength. This confirms the main criticism of the LNM analysis raised in [3] that the LNM values are highly correlated with the node strength of the connectome matrix.

On the other hand, the simulation test using synthetic lesion and connectivity matrices shows that the LNM method can actually detect the network of regions associated with a lesion and a given set of clinical symptoms. LNM values increase for highly interconnected regions and for regions with frequent lesion occurrence. However, this is relative to the node strength of the corresponding region rather than to zero.

The *NullR* sensitivity test results provide another explanation for the concerns raised in [3] regarding the similarity of sensitivity test results across studies. Statistical tests against a zero mean or any horizontal baseline identify the regions in the upper right part of the “LNM-cloud” (see Fig. 1 **f**), thereby detecting the regions with higher node strength. This is a consequence of the linear dependence of the expected LNM value on the node strength and of an additional correlation (a tilt) between a region’s node strength and lesion occurrence (see black dots in **f**). Thus, the effect observed and reported in [3] that the results of the standard sensitivity test appear similar for different clinical cases and are related to the property of the matrix **C** is valid. However, the proposed *NullR* test identifies regions independently of their node strength and, nevertheless, has high overlap with the standard sensitivity test. This is due to the tilted shape of the “LNM-cloud” with respect to the expected value, which is also observed in other clinical cases (see Fig. S14).

One of the main criticisms raised by [4] in their reply to [3] was that the specificity analysis had not been adequately addressed. On the other hand, [3, 20] expressed concerns that the specificity test may not be independent of the sensitivity test. According to the authors, the sensitivity test converges on the node strength of the normative connectome matrix **C**. They therefore argued that the two-sample t-test effectively compares two groups of values related to node strength with different means. Moreover, in [20], where this topic is addressed as a separate study, the authors argue that the shape of the “LNM-cloud” also fully reflects the properties of the principal components of the connectome matrix **C**. Nevertheless, the results of the specificity test with the *NullS* model provided a step towards reconciling the two opposing viewpoints. First, defining *LNM*_*k*_ values as an average, rather than the sum of rows (see Eq. (2) and Supplementary Section S1 and S2) yields the expected values to be *d*_*k*_*/R* and independent of the sample size *S*. This ensures that the distribution of *LNM* values for the groups with (+) and without (−) symptoms has the same expected values. In other words, the baseline values of *LNM* ^(+)^ and *LNM* ^(−)^ are the same. Second, as it is implemented in the *NullR* model, full permutation of the columns of the lesion matrix 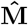, or the rows of the connectivity matrix **C** results in the expected value of LNM being equal to the diagonal. Any deviation in the shape of the “LNM-cloud” comes from properties of the lesion matrix 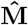 that are specific to the given clinical case (see Supplementary Section S6.4). Furthermore, the regions selected in the standard and *NullS* specificity tests are not biased towards high node strength values. This property also affects the selection of regions in the conjunction map, i.e., the lesion- and symptom-associated network, ensuring that it is not biased by higher node strength. Moreover, a detailed comparison of the *ttest2* and *NullS* results reveals that *ttest2* is susceptible to detecting outliers for small sample sizes, whereas *NullS* is not due to permutation across all subjects (see Supplementary Fig. S13). However, it should be noted that recent works on LNM have not applied *ttest2* for specificity testing but have also relied on permutation of the subjects’ clinical scores [9, 10], which already addresses this issue. Thus, the proper formulation of the specificity test, and thereby the conjunction map, provides symptom-specific results; i.e., we agree with that the methodology behind LNM is sound. Despite this, the choice of the control group (the group without the given clinical symptom) is an important factor that can impact the result of the specificity test.

Our models *NullR* and *NullS* are based on permutation of the lesion matrix, which makes the assumption that the permuted values are independent. This is a simplified assumption with respect to the lesion-regions association, and more conservative null models could be built using information not included in the lesion or connectivity matrices. The frequency of the lesion distribution, joint probability of the lesion occurrence between different brain regions, vascular atlas of the brain, or information about neural connections between brain regions, e.g. obtained from diffusion tensor imaging, could be used to impose additional constraints on the permutation of the lesion and connectivity matrices. Nevertheless, the liberal permutation-based models *NullR* and *NullS* provide significant information for analysis of the sensitivity and specificity test results, shedding light on the factors underlying the debate in recent works on LNM [3, 4, 7, 20].

The simplified formulation of LNM proposed in [3] already provides a significant boost in computational speed. However, performing permutation tests could still substantially reduce the speed of the analysis. The large size of the voxel-wise connectivity matrix poses a particular computational challenge in this regard. For *R* = 2.8 *×* 10^5^ voxels, the connectivity matrix contains approximately *R × R* = 8.17 *×* 10^10^ entries and requires approximately 327 GB of memory in single precision. This makes explicit permutation-based testing computationally expensive. The conclusions and results obtained for the suggested null models are also applicable to the voxel-wise formulation. The analytical expressions derived for *NullS* and *NullR* (Eqs. (4),(5), (6) and (7)) and also the analytical expression for the Zalesky & Cash null model (Eqs. (S33) and (S36)) provide a substantial computational advantage by reducing the computational time by a factor proportional to the number of permutation repetitions. Furthermore, several terms, including node strength, the variance of the connectivity matrix rows (in Eq. (5)) and the variance of a row over its neighbors (in Eq. (S36)) can be pre-computed once for a given dataset. Additionally, Eq. (7) can be evaluated efficiently using column-wise chunked connectivity matrix multiplication. This further reduces the computational and memory requirements.

In summary, the apparent disagreement between the two approaches stems from their consideration of different aspects of the same LNM analysis. The simplified LNM formulation suggested by [3] correctly reveals the dependence of the LNM values on normative connectome node strength, as demonstrated by our analytical derivations. At the same time, the claim that LNM can identify symptom-specific regions, as addressed by [4–6], is also valid when specificity is performed and tested against an appropriate null model. This model provides results without the bias introduced by the normative connectome’s structure. Conjunction maps resulting from the suggested statistical analysis substantially overlap with the results of the standard approach but also identify additional significant regions (red arrow in Fig. 1 *h*). These regions may provide new targets for investigation in future studies.

## Supplementary information

This article has one accompanying Supplementary File.

## Acknowledgements

We thank Prof. Dr. Andreas Horn and Christine Paulus for their valuable scientific input and constructive feedback.

## Declarations

### Ethics approval and consent to participate

N/A

### Consent for publication

N/A

### Data availability

Repositories of data used are cited in the text.

### Materials availability

N/A

## Footnotes

1 Here and throughout the text, the superscript (^∗^) denotes the corresponding permuted variable.

2 The dice coefficient between two sets *A* and *B* is defined as 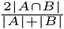.

