## Supplementary Material for "Reconciling Two Sides: Novel Analytical Statistics for Lesion Network Mapping"

#### S1 Definitions

Notations that are used in this report:

1. $R$ is the number of regions.
2. $S$ is the number of separate measurements, which are sometimes referred to as lesions.
3. $\mathbf{C}$ is the group average functional connectivity matrix calculated from the dataset of resting-state fMRI of healthy subjects. The matrix $\mathbf{C}$ is symmetric of size $R \times R$ and is constant for a given dataset.
4. $\mathbf{M}$ is the binary lesion matrix. Every row is a vector $\mathbf{m_s}$ of size $1 \times R$. The entries are 1 if the region is affected by a lesion and 0 if it is not.
5. In [1], a row-normalized lesion matrix is used, i.e., the binary lesion matrix $\mathbf{M}$ is divided by the lesion size $|\mathbf{m_s}|$. We denote this matrix as:

$$\hat{\mathbf{M}}_{ij} = \frac{M_{ij}}{m_i}, \tag{S1}$$

where $m_i = |\mathbf{m_s}|$ is the lesion size, i.e., the number of affected regions in the measurement (lesion) $i$:

$$m_i = \sum_{j=1}^{R} M_{ij}. \tag{S2}$$

Thus, the non-zero elements of the lesion matrix are equal to 1 or $1/|\mathbf{m_s}|$ if several regions are affected by a lesion.

6. $d_k = \sum_{j=1}^{R} C_{jk}$ is the node strength of a region $k$.

According to [1], the lesion network mapping can be written as follows

$$\mathbf{LNM} = \hat{\mathbf{M}} \times \mathbf{C}, \tag{S3}$$

which results in a matrix of size $S \times R$, and every row corresponds to fingerprints in the standard LNM analysis [1]. The actual value used to find the network is defined in [1] as a sum

$$LNM = \sum_{i=1}^{S} \hat{\mathbf{M}} \times \mathbf{C}. \tag{S4}$$

[2] defines it through averaging:

$$LNM = \frac{1}{S} \sum_{s=1}^{S} \mathbf{M} \times \mathbf{C}. \tag{S5}$$

Note that [2] considered the case when one lesion covers one region only ($m_i = 1, i = 1, \ldots, S$). In that case, the lesion matrix remains binary ($\hat{\mathbf{M}} = \mathbf{M}$) and the lesion network mapping can be written as

$$LNM = \sum_{s=1}^{S} \mathbf{M} \times \mathbf{C}. \tag{S6}$$

For our simplified formulation of $LNM$ we use the averaged form, as in Eq. S5, with the lesion matrix $\hat{\mathbf{M}}$:

$$LNM = \frac{1}{S} \sum_{s=1}^{S} \hat{\mathbf{M}} \times \mathbf{C}, \tag{S7}$$

which produces a vector of size $1 \times R$, where each element will be denoted as $LNM_k$. As will be shown below, this definition yields an interpretable expected value for the null distribution independent of the sample size.

We define the mean frequency of a region to be affected across all patient lesions:

$$f_j = \frac{1}{S} \sum_{i=1}^{S} \frac{M_{ij}}{m_i} = \frac{1}{S} \sum_{i=1}^{S} \hat{M}_{ij}. \tag{S8}$$

Then for region $k$ we have:

$$LNM_k = \frac{1}{S}\sum_{i=1}^{S}\frac{1}{m_i}\sum_{j=1}^{R} M_{ij}C_{jk} = \sum_{j=1}^{R}\frac{1}{S}\sum_{i=1}^{S}\frac{M_{ij}}{m_i}C_{jk} = \sum_{j=1}^{R} f_j C_{jk}. \tag{S9}$$

As matrix formulation this reads:

$$LNM = \mathbf{f} \times \mathbf{C}.$$

#### S2 Derivation of Null Models

According to the sensitivity and specificity tests, we derive two distinct null models and null hypotheses, each with its own randomization procedure. Both models treat the values of $LNM$ as the product of a row-normalized lesion matrix $\hat{\mathbf{M}}$ and a symmetric connectivity matrix $\mathbf{C}$ (see Eq. (S7)). Both tests apply randomization to this fixed dataset using physically meaningful permutations of the labels: a permutation of the columns, i.e., the regions, and a permutation of the rows, i.e., the subjects or symptoms.

The two null models are the following:

1. Column permutation or *NullR* model:
   - $\mathbf{H_0}$: The observed assignment of lesion frequencies to the regions is exchangeable.
   - $\mathbf{H_1}$: The values of $LNM_k$ are systematically larger than those produced by random region-label assignment.
   - **Permutation**: We randomly permute the columns of $\hat{\mathbf{M}}$ to shuffle the regions across all patients and obtain a null distribution of the mean values of $LNM$ per region. In this case, $m_i$ is not permuted.
2. Row-split permutation or *NullS* model:
   - $\mathbf{H_0}$: Membership in the (+) and (−) symptom groups is interchangeable.
   - $\mathbf{H_1}$: The difference in means between the two groups $\Delta = LNM^{(+)} - LNM^{(-)}$, is systematically larger than what would be produced by randomly splitting $S = n_+ - n_-$ subjects into the (+) and (−) groups of sizes $n_+$ and $n_-$.
   - **Permutation**: We estimate, region-wise, a null distribution for the difference, $\Delta = LNM^{(+)} - LNM^{(-)}$, by uniformly partitioning the $S$ rows randomly into two groups of sizes $n_+$ and $n_-$. This is equivalent to permuting the rows of the lesion matrix $\hat{\mathbf{M}}$ and splitting it into two matrices sized $n_+$ and $n_-$.

First, we introduce the necessary property of the sum of two vectors under uniform permutation. Then, using this property, we derive the expected value and variance of the null distributions.

##### S2.1 Variance of a sum of two vectors under uniform permutation

Let us consider the following sum

$$\sum_{i=1}^{N} a_i b_i,$$

where $a_i$ and $b_i$ with $i = 1, \ldots, N$ are fixed sequences. Now we consider a uniform random permutation $\pi(i)$ of indices $\{1, \ldots, N\}$ applied to the vector $b$. Then the permuted sum is

$$T^* = \sum_{i=1}^{N} a_i b_{\pi(i)}. \tag{S10}$$

Here and further, we denote a random permutation of a variable with $^*$, i.e., $b_i^* = b_{\pi(i)}$.

The variance of the sum $T^*$ is, by definition,

$$\mathrm{Var}\left(T^*\right) = \sum_i a_i^2 \,\mathrm{Var}\left(b_i^*\right) + \sum_{i \neq j} a_i a_j \,\mathrm{Cov}\left(b_i^*, b_j^*\right). \tag{S11}$$

Consider the sum of the elements of the vector $b$. The permutation $\pi$ does not change the sum, thus its variance is equal to zero

$$\mathrm{Var}\left(\sum_i b_i^*\right) = 0 = \sum_i \mathrm{Var}\left(b_i^*\right) + \sum_{i \neq j} \mathrm{Cov}\left(b_i^*, b_j^*\right). \tag{S12}$$

Since the permutation is uniform, every position $i$ has the same variance and is equal to the variance of the original $N$ values:

$$\mathrm{Var}\left(b_i^*\right) = \sigma_b^2 = \frac{1}{N} \sum_i (b_i - \bar{b})^2, \text{where } \bar{b} = \frac{1}{N} \sum_i b_i. \tag{S13}$$

Moreover, every pair of $i \neq j$ has the same joint distribution, and thus the same covariance: $\mathrm{Cov}\left(b_i^*, b_j^*\right) = \rho_b$. Plugging it into the variance of the sum expression Eq. (S12), we have

$$0 = \sum_i \mathrm{Var}\left(b_i^*\right) + \sum_{i \neq j} \mathrm{Cov}\left(b_i^*, b_j^*\right) = \sum_i \sigma_b^2 + \sum_{i \neq j} \rho_b = \tag{S14}$$

$$= N\sigma_b^2 + 2\frac{N(N-1)}{2}\rho_b, \tag{S15}$$

thus

$$\rho_b = -\frac{\sigma_b^2}{N-1}. \tag{S16}$$

Using this expression, we have for Eq. (S11):

$$\mathrm{Var}\left(T^*\right) = \frac{\sigma_b^2}{N-1}\left((N-1)\sum_i a_i^2 - \sum_{i\neq j} a_i a_j\right).$$

Consider the following square of the sum

$$\left(\sum_i a_i\right)^2 = \sum_i a_i^2 + \sum_{i\neq j} a_i a_j.$$

Thus, we have

$$\begin{aligned}
\mathrm{Var}\left(T^*\right) &= \frac{\sigma_b^2}{N-1}\left((N-1)\sum_i a_i^2 - \left(\sum_i a_i\right)^2 + \sum_i a_i^2\right) = \\
&= \frac{\sigma_b^2}{N-1}\left(N\sum_i a_i^2 - \left(\sum_i a_i\right)^2\right) = \\
&= \sigma_b^2 \frac{N^2}{N-1}\left(\frac{1}{N}\sum_i a_i^2 - \left(\frac{1}{N}\sum_i a_i\right)^2\right) = \\
&= \sigma_b^2 \frac{N^2}{N-1}\frac{1}{N}\sum_i \left(a_i - \bar{a}\right)^2,
\end{aligned}$$

where we used the fact that

$$\mathrm{Var}\left(a\right) = \sigma_a^2 = \frac{1}{N}\sum_i \left(a_i - \bar{a}\right)^2 = \frac{1}{N}\left(\sum_i a_i^2 - N\bar{a}^2\right). \tag{S17}$$

Then, the final expression for the variance of the sum is

$$\mathrm{Var}\left(T^*\right) = \frac{N^2}{N-1}\sigma_a^2\sigma_b^2. \tag{S18}$$

Note that in some programming languages the variance is defined as sample variance with Bessel's correction, i.e., with division by $1/(N-1)$. We denote this variance with "hat" as

$$\hat{\sigma}_a^2 = \frac{1}{N-1}\sum_i \left(a_i - \bar{a}\right)^2. \tag{S19}$$

Then, using the sample variance, we write Eq. (S18) as

$$\mathrm{Var}\left(T^*\right) = (N-1)\hat{\sigma}_a^2\hat{\sigma}_b^2. \tag{S20}$$

##### S2.2 *NullR* model: Column permutation

***Expected value.***

We find the region-wise expected value for the *NullR* distribution using the definition given in Eq. S9

$$\mathbb{E}\left[LNM_k^*\right] = \mathbb{E}\left[\sum_{j=1}^{R} f_j^* C_{jk}\right] = \sum_{j=1}^{R} \mathbb{E}\left[f_j^*\right] C_{jk}, \tag{S21}$$

where $\mathbb{E}\left[f_j^*\right]$ depends on the expected value of the permuted lesion matrix $\hat{\mathbf{M}}^*$. Since uniform column permutation makes every column equally likely, the expected value of the frequency $f_j^*$ is simply the average frequency across all columns:

$$\mathbb{E}\left[f_j^*\right] = \frac{1}{R}\sum_{j=1}^{R} f_j = \frac{1}{SR}\sum_{j=1}^{R}\sum_{i=1}^{S}\frac{M_{ij}^*}{m_i} = \frac{1}{SR}\sum_{i=1}^{S}\frac{1}{m_i}\sum_{j=1}^{R} M_{ij} = \frac{1}{SR}\sum_{i=1}^{S} 1 = \frac{1}{R}, \tag{S22}$$

where we used the fact that the column permutation does not change the value of the lesion sizes $m_i$.

Thus, from Eq. (S21) we get:

$$\mathbb{E}\left[LNM_k^*\right] = \sum_{j=1}^{R} \mathbb{E}\left[f_j^*\right] C_{jk} = \frac{1}{R}\sum_{j=1}^{R} C_{jk} = \frac{d_k}{R}. \tag{S23}$$

This formula shows that the expected value of $LNM$ for a given region $k$ under uniform column permutation is simply the node strength of region $k$ divided by the total number of regions.

**Note:** using a row-normalized lesion matrix $\hat{\mathbf{M}}$ and defining the LNM values as the mean (Eq. (S9) and Eq. (S5)) rather than as the sum (Eq. (S4)) cancels out $S$ in Eq. (S22), making the expected value of LNM (Eq. (S23)) independent of the sample size $S$.

***Variance.***

To find the variance of Eq. (S9), we use Eq. (S18): the permuted vector is $f$, and the constant one is the $k$-th column of $\mathbf{C}$: $C_k := C_{\cdot,k}$; the summation is over the regions. Thus, the variance is

$$\mathrm{Var}\left(LNM_k^*\right) = \frac{R^2}{R-1}\sigma_f^2\sigma_{C_k}^2. \tag{S24}$$

If we use the sample variance definition, then we have

$$\mathrm{Var}\left(LNM_k^*\right) = (R-1)\hat{\sigma}_f^2\hat{\sigma}_{C_k}^2.$$

#### S2.3 *NullS* model: Row-split permutation

To simplify the derivation, we introduce a new variable that indicates to which group the index $s$ belongs

$$I_s = \begin{cases} \frac{1}{n_+}, \text{ if the index } s \in G_+, \\ -\frac{1}{n_-}, \text{ if the index } s \in G_-, \end{cases} \tag{S25}$$

where $G_+$ and $G_-$ are the sets of row indices that belong to each group. Then the difference of the mean values of LNM can be written as

$$\begin{aligned} \Delta_k &= LNM_k^{(+)} - LNM_k^{(-)} = \\ &= \frac{1}{n_+} \sum_{s \in G_+} \hat{\mathbf{M}} \times \mathbf{C} - \frac{1}{n_-} \sum_{s \in G_-} \hat{\mathbf{M}} \times \mathbf{C} = \\ &= \sum_s I_s \cdot LNM_{sk}. \end{aligned} \tag{S26}$$

***Expected value.***

Under a uniform random split each row $s$ has the probability to be in $G_+$:

$$P(I_s = 1/n_+) = \frac{n_+}{S},$$

and to be in $G_-$:

$$P(I_s = -1/n_-) = \frac{n_-}{S}.$$

Thus

$$\mathbb{E}\left[I_s^*\right] = \bar{I} = \frac{1}{n_+}\frac{n_+}{S} - \frac{1}{n_-}\frac{n_-}{S} = \frac{1}{S} - \frac{1}{S} = 0. \tag{S27}$$

Therefore, from Eq. (S26) the expected value for $\Delta$ is

$$\mathbb{E}\left[\Delta_k^*\right] = \mathbb{E}\left[\sum_s I_s^* \cdot LNM_{sk}\right] = \sum_s \mathbb{E}\left[I_s^*\right] \cdot LNM_{sk} = 0. \tag{S28}$$

***Variance.***

As shown in Eq. (S27), the mean of $I$ is zero. Moreover, under permutation, the sum remains unchanged, and the variance of the permuted vector $I_s^*$ is identical to the variance of the original vector $I_s$. Thus, the variance is

$$\begin{aligned} \mathrm{Var}\left(I^*\right) &= \frac{1}{S}\sum_s (I_s - \bar{I})^2 = \frac{1}{S}\sum_s (I_s)^2 = \\ &= \frac{1}{S}\left(n_+\left(\frac{1}{n_+}\right)^2 + n_-\left(-\frac{1}{n_-}\right)^2\right) = \frac{1}{S}\left(\frac{1}{n_+} + \frac{1}{n_-}\right). \end{aligned}$$

Using Eq. (S18), where the permuted variable is $I_s$, the constant vector the $k$-th column of $LNM$: $LNM^{(k)} := LNM_{\cdot,k}$, and summation is over the subjects $S$, we have

$$\mathrm{Var}\left(\Delta_k^*\right) = \frac{S^2}{S-1}\frac{1}{S}\left(\frac{1}{n_+} + \frac{1}{n_-}\right)\sigma^2_{LNM^{(k)}} = \tag{S29}$$

$$= \left(\frac{1}{n_+} + \frac{1}{n_-}\right)\frac{S}{S-1}\sigma^2_{LNM^{(k)}} = \left(\frac{1}{n_+} + \frac{1}{n_-}\right)\hat{\sigma}^2_{LNM^{(k)}}, \tag{S30}$$

where we took into account that the sample variance of the $k$-th column of $LNM$ is

$$\hat{\sigma}^2_{LNM^{(k)}} = \frac{1}{S-1}\sum_s \left(LNM_{sk} - LNM_k\right)^2 = \frac{S}{S-1}\sigma^2_{LNM^{(k)}}.$$

### S3 Zalesky & Cash null model

***Procedure.***

To find the null distribution used in [2], the elements of the lesion matrix $\hat{\mathbf{M}}$ are permuted using the following procedure:

- For every node, define a neighborhood as the nodes with the closest absolute values of node strength (20% of the closest).
- The lesions (affected regions or nodes) can be permuted to any of their neighbors, allowing for overlap. This means that one column of the permuted lesion matrix $\hat{\mathbf{M}}^*$ can contain several elements from different columns (regions) of the lesion matrix $\hat{\mathbf{M}}$.
- If one node has several lesions within a row, the final element of the shuffled matrix $\hat{\mathbf{M}}^*$ is the sum of the overlapping lesions.
- The null distribution is built as the set of the largest $LNM$ values of every permutation.

***Neighbor and transition matrices.***

We denote the number of neighbors as $K = \lfloor 0.2R \rfloor$ and define a binary neighborhood matrix $\mathbf{N}$ of the size $R \times R$ such that $N_{kj} = 1$ if node $j$ is the neighbor of node $k$, i.e., node $j$ is within 20% of the closest to node $k$ in absolute value of the node strength:

$$N_{kj} = \begin{cases} 1, \text{ if } |d_k - d_j| \le \delta_K(k), \\ 0, \text{ otherwise}, \end{cases}$$

where $\delta_{(j)}(k)$ are the ascending sorted values of $\{|d_k - d_j|\}_{j=1}^R$, and the subscript $K$ denotes the $K$ nearest neighbor.

Note that when permuted within a row, the element $k$ is allowed to move to one of the elements $j$ with $N_{kj} = 1$. The neighborhood matrix $\mathbf{N}$ is defined solely by the connectivity matrix $\mathbf{C}$.

We define the following binary matrix $\mathbf{Z}$ of transition:

$$Z_{ik\to j} = \begin{cases} 1, \text{ if element } (i,k) \text{ of } \hat{\mathbf{M}} \text{ moved to } (i,j), \\ 0, \text{ otherwise,} \end{cases}$$

and denote the permuted lesion matrix as $\hat{\mathbf{M}}^*$:

$$\hat{M}^*_{ij} = \sum_{k=1}^{R} \hat{M}_{ik} Z_{ik\to j}.$$

Then, the frequency of a node $j$ after permutation is

$$f^*_j = \frac{1}{S}\sum_{i=1}^{S} \hat{M}^*_{ij} = \frac{1}{S}\sum_{i=1}^{S}\sum_{k=1}^{R} \hat{M}_{ik} Z_{ik\to j}, \tag{S31}$$

and the mean $LNM^*$ value after permutation for region $l$ is

$$LNM^*_l = \sum_{j=1}^{R} f^*_j C_{jl} \tag{S32}$$

***Expected value.***
The probability of a transition from $k$ to $j$ in a row $i$ is

$$\mathbb{P}\left(Z_{ik\to j} = 1\right) = p_{kj} = \frac{N_{kj}}{\sum_j N_{kj}} = \frac{N_{kj}}{K},$$

where we take into account that every region has $K$ neighbors. Thus, the sum of a row is constant and equal to $K$.

Now we search for the expected value and variance for $f^*_j$:

$$\mathbb{E}[f^*_j] = \frac{1}{S}\sum_{i=1}^{S}\sum_{k=1}^{R} \hat{M}_{ik} p_{kj} = \frac{1}{S}\sum_{i=1}^{S}\sum_{k=1}^{R} \hat{M}_{ik}\frac{N_{kj}}{K} = \sum_{k=1}^{R}\frac{N_{kj}}{K}\frac{1}{S}\sum_{i=1}^{S}\hat{M}_{ik} = \sum_{k=1}^{R}\frac{N_{kj}}{K} f_k.$$

Finally, we can find the expected value for the null distribution of $LNM^*_l$ according to [2]:

$$\mathbb{E}\left[LNM^*_l\right] = \sum_{j=1}^{R}\mathbb{E}[f^*_j] C_{jl} = \frac{1}{K}\sum_{j=1}^{R}\sum_{k=1}^{R} N_{kj} f_k C_{jl} = \frac{1}{K}\sum_{k=1}^{R} f_k \sum_{j=1}^{R} N_{kj} C_{jl}. \tag{S33}$$

***Variance.***
We find the variance of $f^*$ by using the formula for the variance of a weighted sum (Eq. (S31), $\hat{\mathbf{M}}$ is constant, and $\mathbf{Z}$ is random) and the fact that $\mathbf{Z}$ is binomial:

$$\mathrm{Var}\left(Z_{ik\to j}\right) = p_{kj}(1-p_{kj}).$$

Thus,

$$\begin{aligned}
\mathrm{Var}\left(f_j^*\right) &= \frac{1}{S^2}\sum_{i=1}^{S}\sum_{k=1}^{R}\hat{M}_{ik}^2\,\mathrm{Var}\left(Z_{ik\to j}\right) = \frac{1}{S^2}\sum_{i=1}^{S}\sum_{k=1}^{R}\hat{M}_{ik}^2 p_{kj}(1-p_{kj}) = \\
&= \frac{1}{K^2}\sum_{k=1}^{R}N_{kj}(K-N_{kj})\frac{1}{S^2}\sum_{i=1}^{S}\hat{M}_{ik}^2 = \frac{1}{K^2}\sum_{k=1}^{R}N_{kj}(K-N_{kj})\tilde{f}_k^{(2)} = \\
&= \frac{1}{K^2}\sum_{k=1}^{R}(K-1)N_{kj}\tilde{f}_k^{(2)} = \frac{K-1}{K^2}\sum_{k=1}^{R}N_{kj}\tilde{f}_k^{(2)},
\end{aligned}$$

where we denote $\tilde{f}_k^{(2)} = 1/S^2\sum_{i=1}^{S}\hat{M}_{ik}^2$ and take into account that $\mathbf{N}$ is a binary matrix ($N_{kj}^2 = N_{kj}$).

With the given permutation procedure, the regions are "competing" since they can share the same neighbors. Thus, the covariance between frequencies after permutation is not zero. To find the covariance of $f^*$ we find the covariance of $\mathbf{Z}$ for two different target nodes $j$ and $j'$ for fixed $(i,k)$, namely

$$\mathrm{Cov}\left(Z_{ik\to j}, Z_{ik\to j'}\right) = -p_{kj}p_{kj'},$$

and the covariance between frequencies is

$$\begin{aligned}
\mathrm{Cov}\left(f_j^*, f_{j'}^*\right) &= \frac{1}{S^2}\,\mathrm{Cov}\left(\sum_{i,k}\hat{M}_{ik}Z_{ik\to j}, \sum_{i',k'}\hat{M}_{i'k'}Z_{i'k'\to j'}\right) \\
&= \frac{1}{S^2}\sum_{i,k}\hat{M}_{ik}^2\,\mathrm{Cov}\left(Z_{ik\to j}, Z_{ik\to j'}\right),
\end{aligned}$$

where we take into account that only the same $(i,k)$ elements of $\mathbf{Z}$ are dependent, and the others are independent (zero). Then

$$\begin{aligned}
\mathrm{Cov}\left(f_j^*, f_{j'}^*\right) &= -\frac{1}{S^2}\sum_{i,k}\hat{M}_{ik}^2 p_{kj}p_{kj'} = -\frac{1}{S^2}\sum_{i,k}\hat{M}_{ik}^2\frac{N_{kj}N_{kj'}}{K^2} = \\
&= -\frac{1}{K^2}\sum_{k=1}^{R}N_{kj}N_{kj'}\frac{1}{S^2}\sum_{i=1}^{S}\hat{M}_{ik}^2 = -\frac{1}{K^2}\sum_{k=1}^{R}N_{kj}N_{kj'}\tilde{f}_k^{(2)}.
\end{aligned}$$

Thus, the variance for the null distribution of $LNM_l^*$ according to [2] is:

$$\begin{aligned}\mathrm{Var}\left(LNM_l^*\right) &= \sum_{j=1}^{R} C_{jl}^2 \,\mathrm{Var}\left(f_j^*\right) + \sum_{j,j',j\neq j'}^{R} C_{jl}C_{j'l}\,\mathrm{Cov}\left(f_j^*, f_{j'}^*\right) = \\ &= \sum_{j=1}^{R} C_{jl}^2 \frac{K-1}{K^2} \sum_{k=1}^{R} N_{kj}\tilde{f}_k^{(2)} - \sum_{j,j',j\neq j'}^{R} C_{jl}C_{j'l}\frac{1}{K^2}\sum_{k=1}^{R} N_{kj}N_{kj'}\tilde{f}_k^{(2)} = \\ &= \frac{1}{K^2}\sum_{k=1}^{R}\tilde{f}_k^{(2)}\left((K-1)\sum_{j=1}^{R} C_{jl}^2 N_{kj} - \sum_{j,j',j\neq j'}^{R} C_{jl}C_{j'l}N_{kj}N_{kj'}\right). \end{aligned} \tag{S34}$$

Define

$$a_j := N_{kj}C_{jl} = (\mathbf{N}\times\mathbf{C})_{kl},$$

where $\mathbf{N}\times\mathbf{C}$ is an effective neighbors connectivity matrix. Then, due to the binary $\mathbf{N}$, we have

$$a_j^2 = N_{kj}^2 C_{jl}^2 = N_{kj}C_{jl}^2.$$

We use the expression

$$\sum_{j,j',j\neq j'} a_j a_{j'} = \left(\sum_j a_j\right)^2 - \sum_j a_j^2.$$

Therefore, the expression in the brackets in Eq. (S34) can be simplified as

$$\begin{aligned}(K-1)\sum_{j=1}^{R} a_j^2 - \sum_{j,j',j\neq j'}^{R} a_j a_{j'} &= (K-1)\sum_{j=1}^{R} a_j^2 - \left(\left(\sum_{j=1}^{R} a_j\right)^2 - \sum_{j=1}^{R} a_j^2\right) = \\ &= K\sum_{j=1}^{R} a_j^2 - \left(\sum_{j=1}^{R} a_j\right)^2 = K\sigma_{a_j}^2, \end{aligned} \tag{S35}$$

where we took into account Eq. (S17).

The final expression for the variance is

$$\mathrm{Var}\left(LNM_l^*\right) = \frac{1}{K^2}\sum_{k=1}^{R}\tilde{f}_k^{(2)} K\,\mathrm{Var}\left((\mathbf{N}\times\mathbf{C})_{kl}\right), \tag{S36}$$

where the variance is calculated for every row $k$ only over its neighbors, which solely depends on the connectivity matrix $\mathbf{C}$ and is fixed for a given normative dataset.

***Threshold value of the null distribution.***

In [2] the null distribution is constructed from the largest $LNM$ values in each permutation round. Thus, we can approximate the null distribution by taking the distribution of the nodes (regions) with the largest expected value and variance. The exact formula for the cumulative distribution function (CDF) for the distribution of the largest value in each permutation is the product of the individual CDFs of each region:

$$\Phi_Y(y) = \prod_{i=1}^{R} \Phi\left(\frac{y-\mu_i}{\sigma_i}\right), \tag{S37}$$

where $\mu_i = \mathbb{E}\left[LNM_i^*\right]$, $\sigma_i = \sqrt{\mathrm{Var}\left(LNM_i^*\right)}$, $\Phi$ is the CDF of the normal distribution, $Y = \max(LNM_i^*), i = 1, \ldots, R$.

Since the expected values of $LNM$ depend on the node strengths of the region, the z-scores in Eq. (S37) for regions with lower node strength, i.e., lower $\mu_i$, will be high. Accordingly, the CDF values will be close to 1. Thus, only the regions with high expected values will contribute to the final CDF $\Phi_Y$.

In practice, we calculate the expected values and variance for all nodes using Eqs. (S33) and (S36), then select the $k$ regions with the highest scores $\mathbb{E}\left[LNM_i^*\right] + \Phi^{-1}(p)\,\mathrm{Var}\left(LNM_i^*\right)$. Here $\Phi^{-1}$ is the inverse CDF of the normal distribution and $p$ is the target percentile. Then, by finding the values of $\Phi_Y$ as

$$\Phi_Y(y) = \prod_{i=1}^{k} \Phi\left(\frac{y-\mu_i}{\sigma_i}\right), \tag{S38}$$

within the range of $\mathbb{E}\left[LNM_x^*\right] \pm 3\,\mathrm{Var}\left(LNM_x^*\right)$, where $x$ is the index of the region with the largest $\mathbb{E}\left[LNM_x^*\right]$, we find the threshold value that corresponds to the given percentile $p$.

Taking $k = 15$ already gives a good approximation of the right side of the Zalesky & Cash null distribution as shown in Fig. S3 in the next section. We will denote this analytical expression of the Zalesky & Cash null distribution threshold value as $LNM_{zc}$.

***Calculation.***

The neighborhood matrix $\mathbf{N}$ as well as the effective neighborhood connectivity matrix $\mathbf{N} \times \mathbf{C}$ are determined entirely by the averaged connectivity matrix $\mathbf{C}$ and can be precomputed once for a given dataset and neighborhood size $K$. The variance in Eq. (S36) is also entirely defined by the connectivity matrix $\mathbf{C}$ and can be computed in advance.

Therefore, for a given lesion matrix $\mathbf{M}$, only two quantities need to be computed. The first is the expected value $\mathbb{E}[f_j^*]$, which is the mean frequency of the regions neighboring region $j$. It is computed as `Efj = sum(f.*N(:,j),1)`. The second is $\tilde{f}_k^{(2)} = \frac{1}{S^2}\sum_{i=1}^{S}\hat{\mathbf{M}}_{ik}^2$, which is computed as `sum(hM.^2,1)/S^2`.

Thus, computing the null distribution using the analytical formulae in Eqs. (S33) and (S36) requires substantially less computational effort than the permutation-based approach, especially when the connectivity matrix is defined for voxels. Furthermore, to find the threshold $LNM_{zc}$, we only need to calculate these values for several nodes, i.e., the nodes with the highest node strength.

#### S4 Verification of the analytical expressions for the null distributions by permutation

We tested all analytical expressions for the null distributions with the outcome of the corresponding permutations. We used either the synthetic connectivity and lesion matrices (see Section S5) or the precomputed group connectivity matrix $\mathbf{C}$ for the GSP1000 (Brain Genomics Superstruct Project 1000 [3]) dataset (see Section S6) as the connectivity matrix. The lesion matrix was taken from the aphasia recovery dataset [4], which was obtained from the GitHub repository [5] of [1].

To test our *NullR* model, we performed the column permutation for the entire lesion matrix $\hat{\mathbf{M}}$ 5000 times. Then, we found an approximation of the normal distribution from the normalized empirical histogram and compared it with the theoretical normal distribution, which was calculated using the expressions Eqs. (S23) and (S24). An example result is shown in Fig. S1.

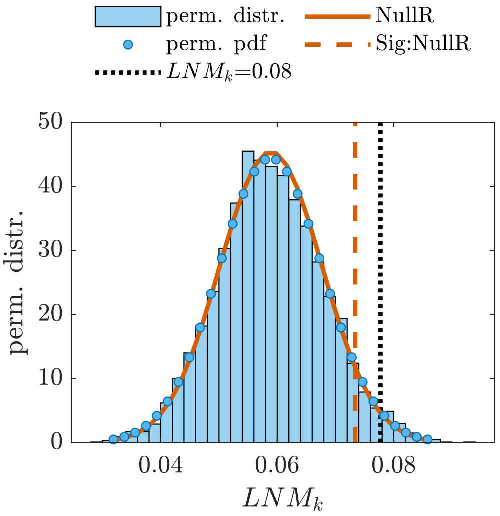


**Fig. S1** Verification of the theoretical expressions for the *NullR* model. The columns of the clinical lesion matrix $\mathbf{M}$ [4] were permuted. The group connectivity matrix $\mathbf{C}$ was taken from GSP1000 [3]. Results are shown for region $k = 718$. The empirical probability density function (pdf) approximated as normal distribution (blue circles), derived from the normalized histogram of the permutation results (blue bars), is compared with the theoretical *NullR* distribution (orange solid line) calculated using Eqs. (S23) and (S24). The vertical orange dashed line denotes the $p = 0.05$ significance threshold of the *NullR* distribution, while the actual value of the $LNM_k$ (vertical black dotted line) falls within the significance range.

To verify the analytical expressions for the *NullS* model, we performed 5000 row permutations of the lesion matrix $\hat{\mathbf{M}}$. After each permutation, we relabeled the first $n_+$ rows as the members of the group with clinical symptoms and the last $n_-$ rows as the group without symptoms. Then, we calculated the LNM values for each group and found the difference $\Delta^* = LNM^{(+)*} - LNM^{(-)*}$ for each permutation round.

The comparison of the permutation results and the theoretical *NullS* distribution is shown in Fig. S2

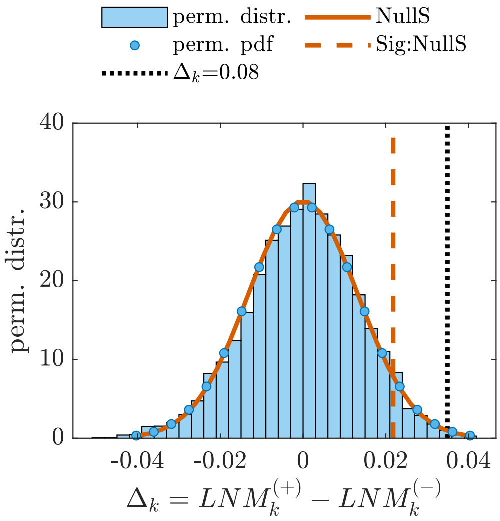


**Fig. S2** Verification of the theoretical expressions for the *NullS* model. The columns of the clinical lesion matrix **M** [4] were permuted. The first $n_+ = 188$ rows were labeled as having symptoms and the last $n_- = 30$ rows as not having symptoms. The group connectivity matrix **C** was taken from GSP1000 [3]. The empirical probability density function (pdf) approximated as normal distribution (blue circles), derived from the normalized histogram of the permutation results (blue bars) for the difference $\Delta$, is compared with the theoretical *NullS* distribution (orange solid line) calculated using Eqs. (S28) and (S30). The vertical orange dashed line denotes the $p = 0.05$ significance threshold of the *NullS* distribution, while the actual value of $\Delta_k$ (vertical black dotted line) falls within the significance range.

We tested the theoretical expressions Eqs. (S33) and (S36) for approximating the Zalesky & Cash null distribution using the procedure described above (S3). We performed the within-neighbor-only permutation procedure described above in Section S3 5000 times, taking the largest value of $LNM$ in every permutation. Using the expressions, we identified $k = 15$ regions with the highest scores $\mathbb{E}\left[LNM_k^*\right] + \Phi^{-1}(0.95)\,\mathrm{Var}\left(LNM_k^*\right)$ and calculated the product of the cumulative distribution functions (see Eq. (S38)). We compared this theoretical *NullZC* distribution with the empirical one. The results are shown in Fig. S3.

The empirical null distribution (the blue bars in Fig. S3) appears to be slightly right-skewed. This is due to the fact that in every permutation round, the largest $LNM$ value is taken. Specifically, taking the maximum of two random variables with closely lying normal distributions causes the left side of the empirical null distribution to be ”shorter” than the right one. However, the right side is defined by the normal distribution of the regions with the largest expected value and variance: compare the right side of the empirical null distribution (blue bars) in Fig. S3 with the theoretically obtained null distribution (solid orange line). A good approximation of the right side of the Zalesky & Cash null distribution also allows one to approximate the threshold value $LNM_{th}$: compare $LNM_{th}$ obtained from the empirical null distribution (the vertical black dotted line) and that obtained using the theoretical normal distribution calculated by Eqs. (S33) and (S36) (the vertical dashed orange line).

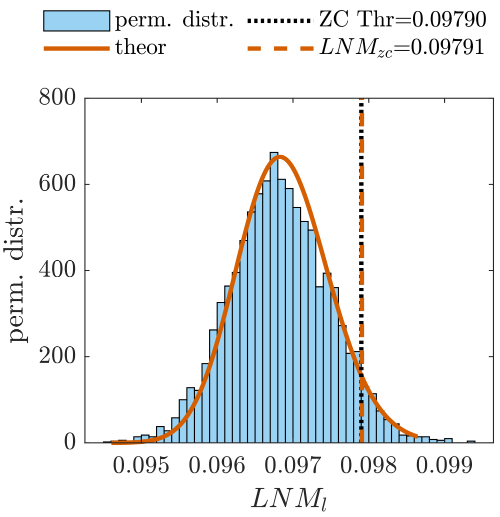


**Fig. S3** Verification of the theoretical expressions for the *NullZC* model distribution using the clinical lesion matrix **M** [4] and the group connectivity matrix **C** for the GSP1000 dataset [3]. A within-neighbor permutation procedure (see Section S3) was performed. The normalized histogram of the permutation results (blue bars) is compared with the theoretical *NullZC* distribution (orange solid line) calculated using Eqs. (S33), (S36) and (S38) for $k = 15$. The empirical threshold (black dotted line) is very close to the analytical threshold (vertical orange dashed line).

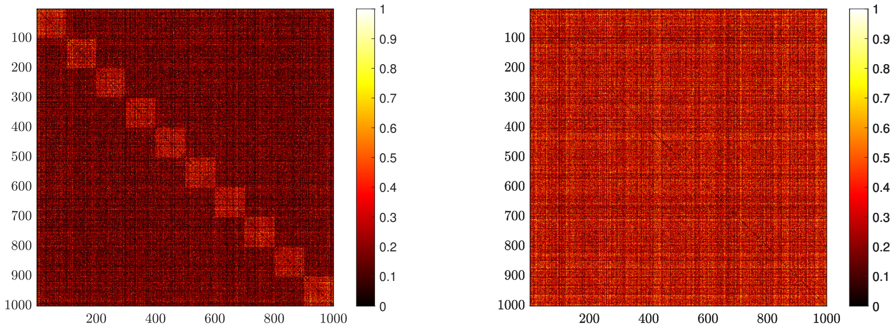


**Fig. S4** Examples of the synthetic log-normal distributed connectivity maps. Left panel: An example with $B_{inter} = 0.3$, $B_{intra} = 0.6$. Connectivity between areas within a target module is higher than between other regions. Right panel: An example with $B_{inter} = 0.3$, $B_{intra} = 0.3$, i.e., a similar probability of being connected with any region.

### S5 Simulation test based on synthetic connectivity and lesion matrices

To test the null models, we used the synthetic connectivity matrix from [2] generated by a degree-controlled stochastic block model. This model produces a unidirectional, weighted symmetric matrix with an approximately log-normal distribution. Please see the supplementary materials of [2] for details on the generation of the synthetic connectivity matrix.

The $R = 1000$ regions were assigned into $B = 10$ equally sized modules, each comprising $B_n = 100$ regions using labeling $z_i \in (1, \ldots, B)$. The connections between regions were defined by a block probability matrix $p$, such that the probability of connections between regions within a module ($p_{ij} = B_{intra}, z_i = z_j$) was different (equal or higher) than the probability of connections between regions from other modules ($p_{ij} = B_{inter}, z_i \neq z_j$). For all simulations, $B_{inter} = 0.3$ was kept constant, and depending on the test, $B_{intra}$ was varied from 0.3 to 0.6. The parameter that controls dispersion around the mean was set to $k = 10$. For visual simplicity, the regions within the same modules were placed near each other in the matrix. Fig. S4 shows two examples of connectivity matrices.

Synthetic lesion sets were generated in the same way as in [2]. The parameter $\alpha$ defined the prioritization of lesion locations (see Fig. S5): if $\alpha = 0$, lesions occur with equal probability across all brain regions; if $\alpha = 1$, all lesions occur within the regions of the target module. If $0 < \alpha < 1$, $\alpha K$ lesions occur within the regions of the target module, while the remaining $(1 - \alpha)K$ lesions are distributed uniformly across the whole brain.

The main goal of the test is therefore to determine whether the method can detect the target module when lesions associated with a given symptom tend to occur within the target region. We investigated three cases: (1) the effect of varying $\alpha$ when $B_{intra} > B_{inter}$; (2) the effect of varying $\alpha$ when $B_{intra} = B_{inter}$; and (3) the effect of changing $B_{intra}$ while keeping $\alpha \neq 0$ fixed. $B_{inter}$ remains constant at 0.3.

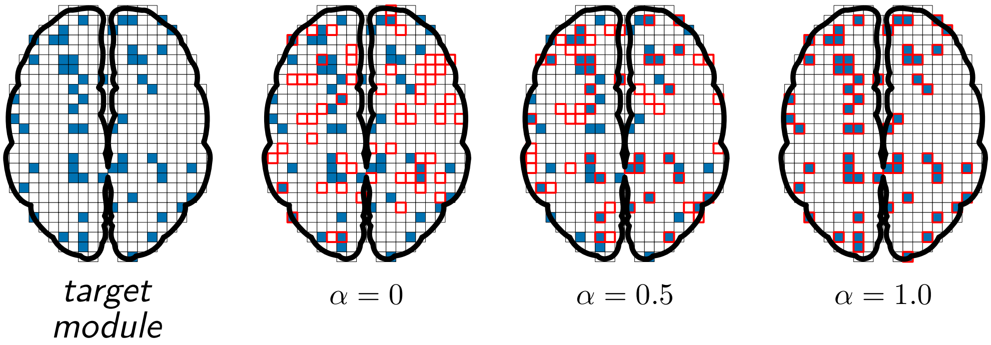


**Fig. S5** Schematic explanation of the synthetic lesions generation. Regions of the target module that are more strongly connected to each other are marked as filled blue squares, while lesions are marked as red squares. When $\alpha = 0$, the lesions occur randomly across the entire brain. When $\alpha = 1$, all lesions occur within the target module regions. When $\alpha = 0.5$, half of the lesions occur within the target module regions.

Fig. S6 shows examples for $B_{intra} = 0.6$ and various values of $\alpha$. An example of the connectivity matrix in this case is shown in Fig. S4 (left). Increasing $\alpha$ causes the LNM values to deviate more from the expected value, i.e., the mean node strength $d/R$. The suggested statistical test can identify regions of the target module, even for small values of $\alpha$.

If the regions of the target module are not topologically distinguishable, i.e., if the inter- and intra-module coupling strengths are equally probable, the connectivity matrix will resemble that shown in Fig. S4 (right). In this case, the LNM values do not differ across regions and modules, irrespective of the value of $\alpha$, which defines the probability that the lesion is within the target module (see Fig. S7). Furthermore, the LNM values are defined by the region's node strength (see Fig. S7, middle panels).

To determine the impact of the intra-module coupling strength on the target module, we kept the values of $\alpha$ and $B_{inter}$ constant while varying $B_{intra}$ (see Fig. S8). Increasing $B_{intra}$ has the same effect as varying $\alpha$: the greater the intra-module connectivity compared to $B_{inter}$, the greater the deviation of the LNM values from the expected value. Once again, the suggested statistical test detects all the regions of the target module as significant.

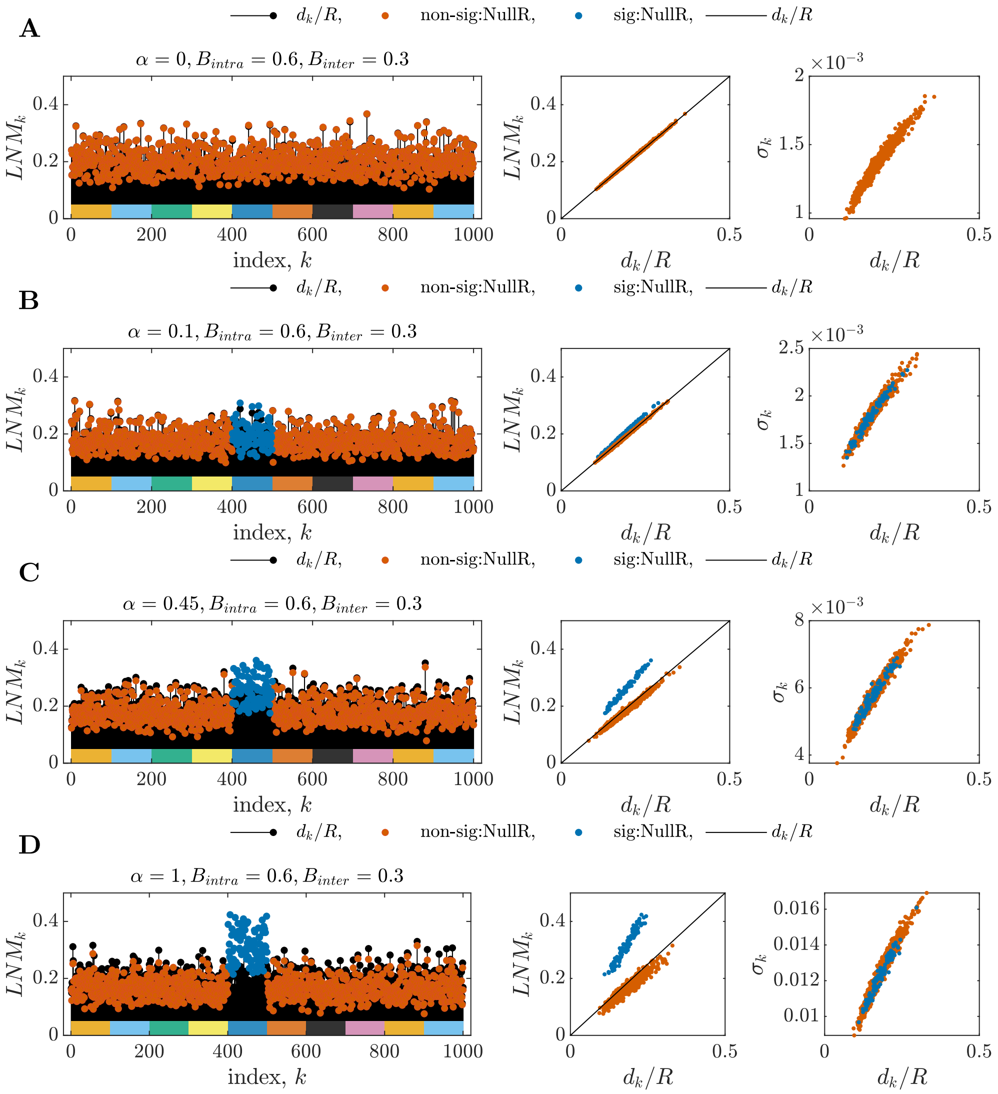


**Fig. S6** Results of the LNM estimation using synthetic connectivity and lesion matrices. The coupling parameters are $B_{inter} = 0.3$ and $B_{intra} = 0.6$. Left panels show the LNM values for all $R = 1000$ regions divided into 10 equally large modules. The black stems represent the expected values of LNM, $d_k/R$. LNM values are plotted by blue dots if they are significant with respect to *NullR* and by orange dots if they are not significant. Middle panels show LNM values plotted against the mean node strength $d/R$. Right panels depict the values of the variance against $d/R$ for every region $k$. (A) $\alpha = 0$. (B) $\alpha = 0.1$. (C) $\alpha = 0.45$. (D) $\alpha = 1.0$.

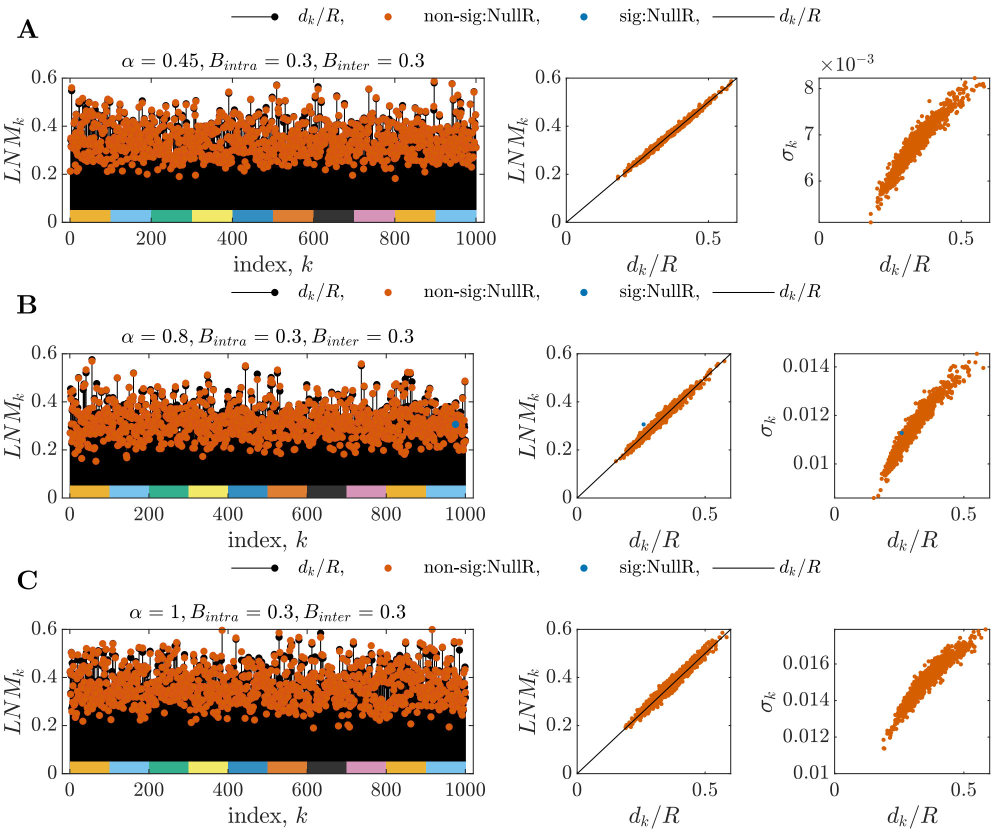


**Fig. S7** Results of the LNM estimation using synthetic connectivity and lesion matrices. The case with equal probability of inter- and intra-module connectivity is shown here: $B_{inter} = 0.3$ and $B_{intra} = 0.3$. (A) $\alpha = 0.45$, (B) $\alpha = 0.8$, (C) $\alpha = 1.0$. All notations are the same as in Fig. S6.

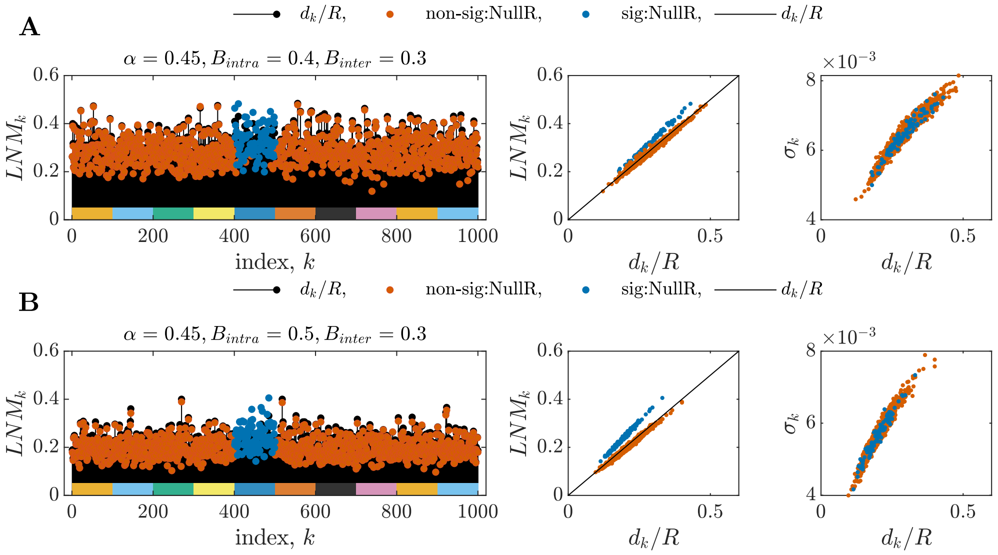


**Fig. S8** Results of the LNM estimation using synthetic connectivity and lesion matrices. The cases with fixed $\alpha = 0.45$ and $B_{inter} = 0.3$, and different (A) $B_{intra} = 0.4$ and (B) $B_{intra} = 0.5$ are presented here. All notations are the same as in Fig. S6.

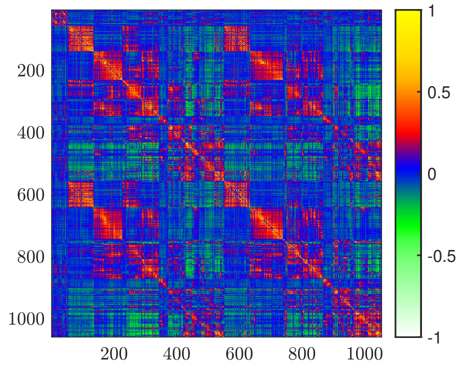


**Fig. S9** The group-averaged and z-scored connectivity matrix from GSP1000 (Brain Genomics Superstruct Project 1000 [3]).

### S6 Details of validation with clinical dataset

#### S6.1 Description of the clinical dataset

We used the connectivity matrix available from the GitHub repository [5] of van den Heuvel and colleagues [1] for the calculations. The averaged connectivity matrix was calculated from the publicly available resting-state functional magnetic resonance imaging dataset (rsfMRI) GSP1000 [3]. The connectivity matrix was mapped to Yeo-Schaefer1000 cortical [6] and Melbourne54 [7] subcortical regions. Therefore, the size of the connectivity matrix $\mathbf{C}$ is $1054 \times 1054$. The connectivity matrix used in the analysis contained the Fisher r-to-z transformed correlation values for each region (see Fig. S9).

The clinical dataset was also taken from the GitHub repository [5] of [1] and from the GitHub repository [8] of [2], which contains lesion matrices for several clinical cases mapped to 1054 cortical and subcortical regions. We used the Aphasia Recovery Cohort (ARC) dataset from [4]. The total number of subjects included in the analysis was 218. The cut-off was set at 93.5 WAB-AQ score, resulting in $n_+ = 188$ subjects with and $n_- = 30$ subjects without aphasia symptoms. The lesion matrix for the aphasia recovery cohort used in the analysis is shown in Fig. S10.

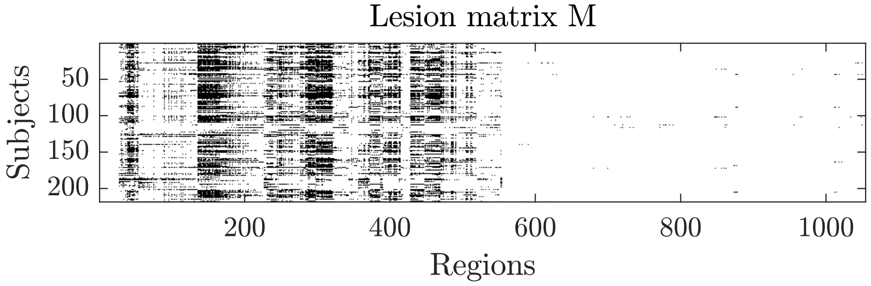


**Fig. S10** The lesion matrix from the aphasia recovery cohort [4].

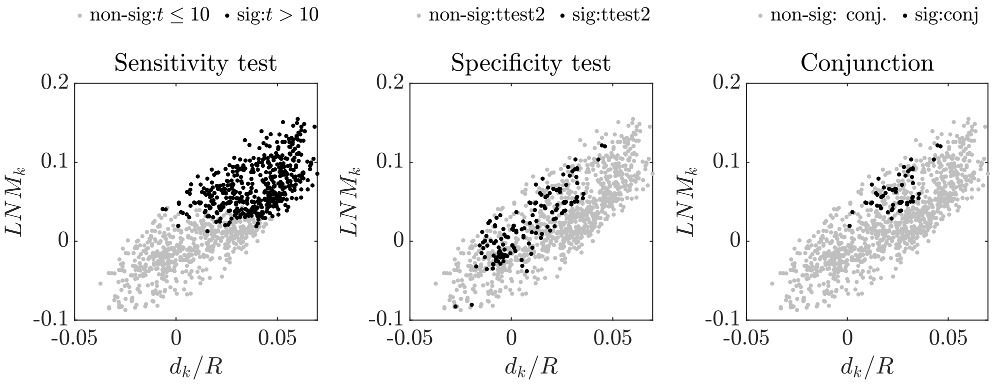


**Fig. S11** Results for the aphasia recovery dataset calculated using standard methods applied to a simplified formulation. Black and gray dots represent regions significant and non-significant according to the corresponding test. Left: sensitivity test performed by applying a region-wise one-sample t-test with zero mean. The threshold is taken as $t > 10$. Middle: specificity test performed by applying a region-wise two-sample t-test on the groups with and without aphasia symptoms. Right: conjunction map constructed by selecting the regions that were significant in the specificity and sensitivity tests.

#### S6.2 Standard LNM analysis

To make the comparison consistent, we performed the standard LNM analysis pipeline using the **LNM** matrix given in Eq. (S7). The rows of the resulting **LNM** matrix are equivalent to the fingerprints used in the standard LNM analysis [1]. The results are shown in Fig. S11.

Unlike in the synthetic case, the LNM values for the real clinical lesion matrix and normative connectivity matrix case are not separated, but form an "LNM cloud" (in all panels in Fig. S11). These clouds can also be found in Fig. 4 in [1], but on different scales.

##### *Sensitivity test*

The standard LNM sensitivity test identifies regions with higher LNM values as significant (black dots in the left panel of Fig. S11). Due to Eq. (S23), those regions also have high node strength; thus, the result of the sensitivity test is biased towards higher node strength.

##### *Specificity test*

We performed the specificity test using a region-wise (column-wise) two-sample t-test with LNM matrices for the group with ($G_+$) and without ($G_-$) symptoms. The results are shown in the middle panel of Fig. S11. As one can see, the regions that are significant according to the two-sample t-test are not biased towards higher or lower node strength and have intermediate values.

##### *Conjunction map*

The conjunction map is constructed by identifying the regions that are significant in both the sensitivity and the specificity tests. The results are shown in the right panel of Fig. S11.

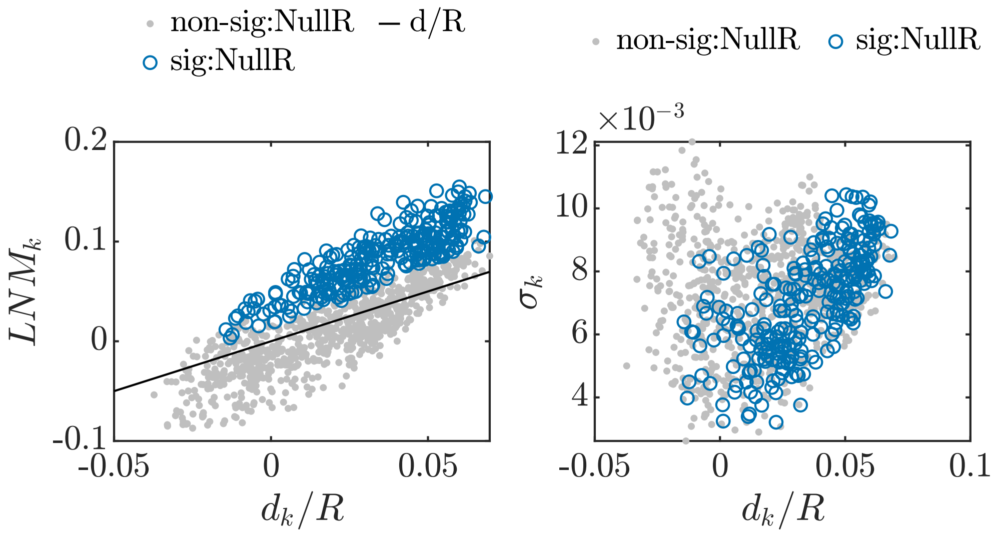


**Fig. S12** Results for the aphasia recovery dataset using the *NullR* test. Left: The significant regions depicted on the $(LNM, d/R)$ plane. The blue circles represent regions that were significant in the NullR test, while the grey dots represent regions that were not significant. The solid black line shows the expected value. Right: The analytically estimated standard deviation of the regions vs $d/R$.

It can be seen that the result of the standard conjunction map can be represented as a subset of the regions significant in the specificity test with values higher than an approximate horizontal threshold defined by the t-value in the sensitivity test. Consequently, the conjunction map results do not contain the regions with high node degree: the significant regions in the upper-right corner in the sensitivity test results (left panel in Fig. S11) are absent from the conjunction map results (right panel in Fig. S11). Thus, the result of the standard conjunction map is not biased towards hub regions, as is the case for the result of the specificity test.

#### S6.3 Suggested statistical method

##### *Results for NullR model*

The result of applying the *NullR* statistical test to the aphasia recovery dataset is presented in Fig. S12. In comparison to the standard sensitivity test, the *NullR* model does not just select regions in the upper part of the "LNM-cloud" (above a horizontal threshold), but rather regions with LNM values above the expected value (i.e., above the diagonal, solid black line in the left panel of Fig. S12). However, the border between the set of significant and not significant regions is not sharp, but rather an overlapping border defined by the individual variance of the regions.

The standard deviation $\sigma_k$ of the regions derived from the real dataset differs from that observed in the synthetic dataset (e.g., compare standard deviation plots in Fig. S6 and Fig. S12). The relationship between the standard deviation and the node strength is not linear. Moreover, the regions that are significant according to *NullR* have a wide range of $\sigma_k$ values.

##### ***Results for NullS model***

*NullS* is calculated using Eqs. (S28) and (S30). Fig. S13 shows the comparison of the results of the two-sample t-test (left panels) and *NullS* (right panels). The upper panels correspond to the results of the group with clinical symptoms and the lower ones to the group without symptoms. As one can see, the two-sample t-test detects outliers (black dots in the upper part of the bottom-left panel in Fig. S13) as significant, whereas *NullS* does not.

Moreover, a comparison of the upper and lower panels in Fig. S13 explains which regions are considered significant in the specificity tests: those with high LNM values in the upper panels and lower values in the lower panels. In this context, it is important to use the correct definition of LNM (as the average, Eq.(S9)) to ensure the same expected mean values for the groups being compared.

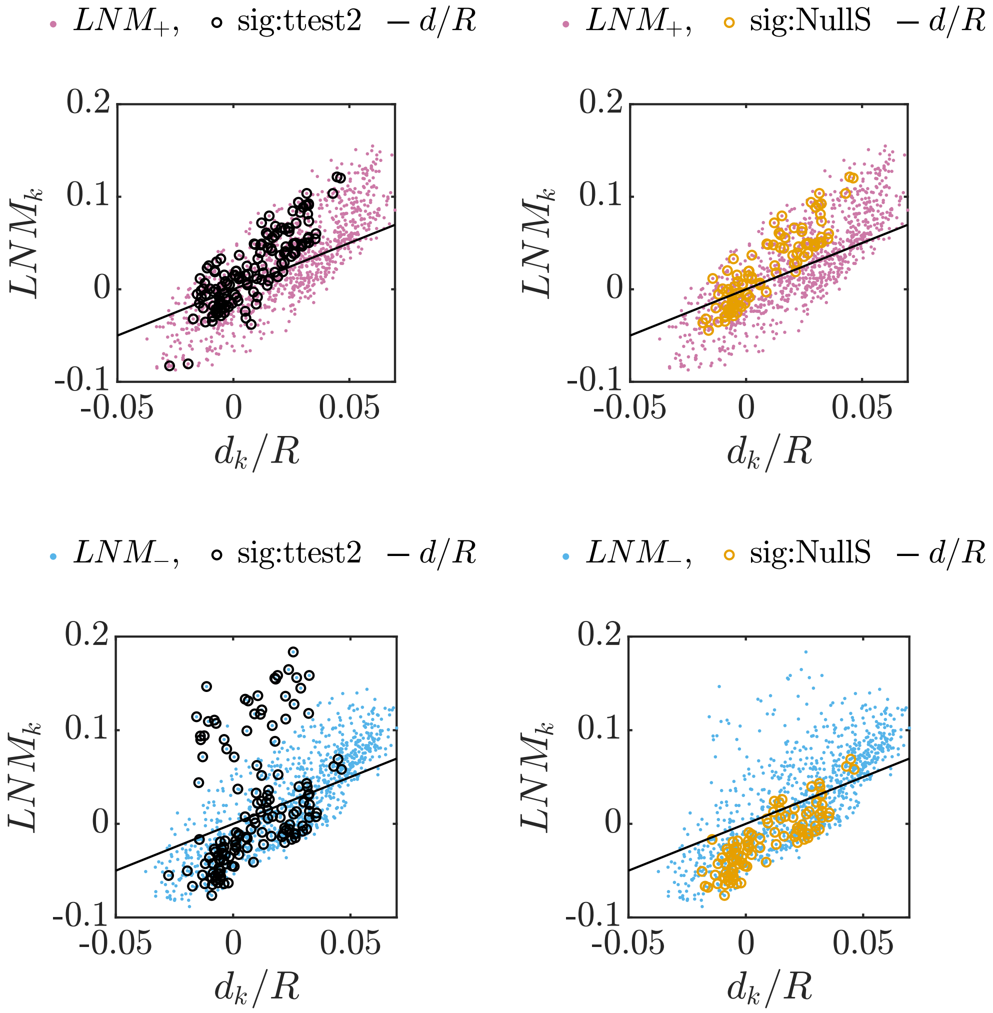


**Fig. S13** Results of the comparison of the two-sample t-test (*ttest2*) and *NullS*. Upper left: *ttest2* results for the group with symptoms. Black circles represent regions significant according to *ttest2*. Lower left: *ttest2* results for the group without symptoms. Black circles represent regions significant according to *ttest2*. Upper right: *NullS* results for the group with symptoms. Orange circles represent regions significant according to *NullS*. Lower right: *NullS* results for the group without symptoms. Orange circles represent regions significant according to *NullS*.

#### S6.4 The shape of the "LNM-cloud"

The "LNM-cloud" for the aphasia recovery dataset has a shape that can be represented with two main principal components. The first principle component is slightly tilted towards the diagonal line, i.e., to the expected value. This shape of the "LNM-cloud" does not originate from the high dimensionality of the connectome matrix $\mathbf{C}$, but rather from the properties of the lesion matrix $\mathbf{M}$. Indeed, the *NullR* model is constructed by full permutation of the columns of the lesion matrix $\mathbf{M}$, which is also equivalent to permuting the rows of the matrix $\mathbf{C}$. Under the full permutation of the columns, the expected value of LNM is the diagonal line. However, the actual observed "LNM-cloud" has a tilted shape, which can only originate from an association between the frequency of the lesion occurring in a given region $f$ and its node strength $d$, i.e., the information obtained from the lesion matrix.

In other words, from Eq. (S9) we see that the LNM value for a given region is the $f$-weighted sum of the columns of $\mathbf{C}$. The expected value of this sum under full permutation of the $f$ vector or the rows of the $\mathbf{C}$ matrix is $d/R$. However, if we observe an "LNM-cloud" with an angle between its principal component and the diagonal line ($d/R$), this comes from the observed distribution of the $f$ vector and arises from the fact that lesions tend to occur more frequently in regions that are positively connected to regions with high node strength and negatively connected to regions with low node strength. Thus, the shape of the "LNM-cloud" reflects the properties of the lesion matrix observed for a given clinical case.

We observe different shapes of the "LNM-cloud" for different clinical cases, as shown in Fig. S14. The shape of the "LNM-cloud" for addiction **a** and insomnia **b** is similar to that of the aphasia recovery results. However, the other cases differ in shape and direction of the main principle components.

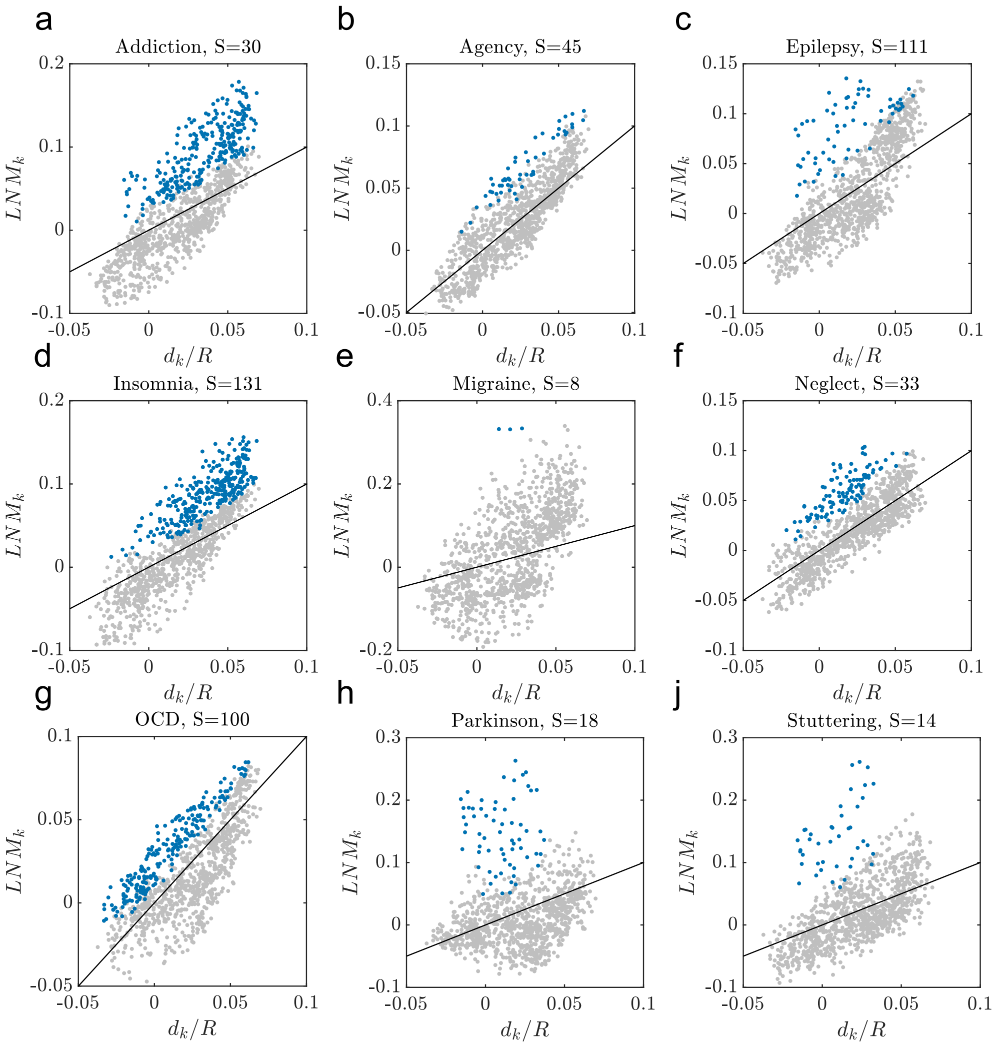


**Fig. S14** The results of *NullR* for different clinical cases. The regions significant according to *NullR* are marked with blue dots, non-significant ones with grey ones. The clinical cases: **a** addition [9], **b** disrupted agency [10], **c** epilepsy [11], **d** insomnia [12], **e** migraine [13], **f** neglect [12], **g** obsessive-compulsive disorder [14], **h** Parkinson's disease [15], **j** stuttering [16].
